# Recovery of small plasmid sequences via Oxford Nanopore sequencing

**DOI:** 10.1101/2021.02.21.432182

**Authors:** Ryan R. Wick, Louise M. Judd, Kelly L. Wyres, Kathryn E. Holt

## Abstract

Oxford Nanopore Technologies (ONT) sequencing platforms currently offer two approaches to whole-genome native-DNA library preparation: ligation and rapid. In this study, we compared these two approaches for bacterial whole-genome sequencing, with a specific aim of assessing their ability to recover small plasmid sequences. To do so, we sequenced DNA from seven plasmid-rich bacterial isolates in three different ways: ONT ligation, ONT rapid and Illumina. Using the Illumina read depths to approximate true plasmid abundance, we found that small plasmids (<20 kbp) were underrepresented in ONT ligation read sets (by a mean factor of ∼4) but were not underrepresented in ONT rapid read sets. This effect correlated with plasmid size, with the smallest plasmids being the most underrepresented in ONT ligation read sets. We also found lower rates of chimeric reads in the rapid read sets relative to ligation read sets. These results show that when small plasmid recovery is important, ONT rapid library preparations are preferable to ligation-based protocols.

**Impact statement:** Researchers who use Oxford Nanopore Technologies (ONT) platforms to sequence bacterial genomes can currently choose from two library preparation methods. The first is a ligation-based approach, which uses ligase to attach sequencing adapters to the ends of DNA molecules. The second is a rapid approach, which uses a transposase enzyme to cleave DNA and attach adapters in a single step. There are advantages to each preparation, for example ligation can produce better yields but rapid is a simpler procedure. Our study reveals another advantage of rapid preparations: they are more effective at sequencing small plasmids. We show that sequencing of ligation-based libraries yields fewer reads derived from small plasmids, making such plasmids harder to detect in bacterial genomes. Since small plasmids can contain clinically relevant genes, including antimicrobial resistance (AMR) or virulence determinants, their exclusion could lead to unreliable conclusions that have serious consequences for AMR surveillance and prediction. We therefore recommend that researchers performing ONT-only sequencing of bacterial genomes should consider using rapid preparations whenever small plasmid recovery is important.

**Data summary:** Supplementary figures, tables, data and code can be found at: github.com/rrwick/Small-plasmid-Nanopore

## Introduction

Plasmids are extra-chromosomal pieces of DNA present in many bacterial genomes^1,2^. While smaller than the chromosome, they are important genomic components that confer key phenotypic traits (including virulence and antimicrobial resistance, AMR) and contribute to gene flow within/between species. Most plasmids are circular, though linear plasmids also exist^3^, and for some bacterial species it is not uncommon to find multiple plasmids in a single genome^4^. Plasmids come in a broad range of sizes, from <1 kbp to >300 kbp^5,6^. For simplicity, here we categorise plasmids as ‘small’ or ‘large’ using a threshold of 20 kbp^7^. Both small and large plasmids can carry clinically relevant genes, such as virulence or AMR genes^8–10^. Unlike many large plasmids, small plasmids do not encode their own conjugative transfer systems, but many do appear to be readily mobilizable between host cells, and are therefore similarly important for the transfer of genetic material and the spread of AMR^11^.

Short-read sequencing (e.g. Illumina platforms) of bacterial genomes can typically only produce fragmented draft assemblies^12^. It is difficult to reconstruct plasmid sequences from short-read assemblies^13^, and this impedes the ability to accurately monitor the spread of key genes in bacterial populations^14,15^. In contrast, long-read sequencing (e.g. Pacific Biosciences and Oxford Nanopore Technologies platforms) allows for complete bacterial genome assembly, with each chromosome or plasmid assembled into a single contig^16,17^. Oxford Nanopore Technologies (ONT) long-read platforms are especially well suited to bacterial genomics, as they are cost-effective and allow for easy multiplexing of samples, enabling sufficient data generation for simultaneous completion of 12 or more genomes at a cost of <100 USD each^18–20^. There are many genome assembly tools appropriate for use with ONT reads, some of which also use Illumina (short) reads for what is known as hybrid assembly^12,21^. Other tools operate on ONT reads alone, an approach we will call long-read-only assembly^22–24^.

While ONT sequencing offers many advantages in bacterial genomics, during a number of sequencing projects^18,25–30^, we have anecdotally observed that small plasmids are frequently absent from ONT long-read-only assemblies. This can have serious downstream consequences, e.g. if a small plasmid containing an AMR determinant is missing from the assembly, this can result in an incorrect antimicrobial susceptibility prediction (a failure to detect resistance, known as a ‘very major error’ as it can lead to prescribing an ineffective antimicrobial^31^). We hypothesised that this problem was due in part to our use of the ONT ligation-based library preparation (SQK-LSK109 kit). During extraction, DNA is incidentally fragmented, typically resulting in fragments ∼10 kbp in size^32^ onto which barcode and adapter sequences are attached via blunt-end ligation during library preparation. Large plasmids are likely to be fragmented into one or more linear pieces, but small plasmids may avoid fragmentation and remain circular. As such, the small plasmids will have no blunt ends, will not have any adapter ligated and will thus be unavailable for sequencing (**Figure 1**). Upon sequencing of the library, this leads to underrepresentation of small plasmids in the resulting ONT read set, which may in turn cause the assembler to fail to produce contigs for them^24^. Deliberately increasing fragmentation before DNA preparation could mitigate this effect by creating smaller fragments, however this would result in shorter read lengths, which negates the benefits of long-read sequencing and can negatively impact the assembly contiguity^24,33^.

**Figure 1:**
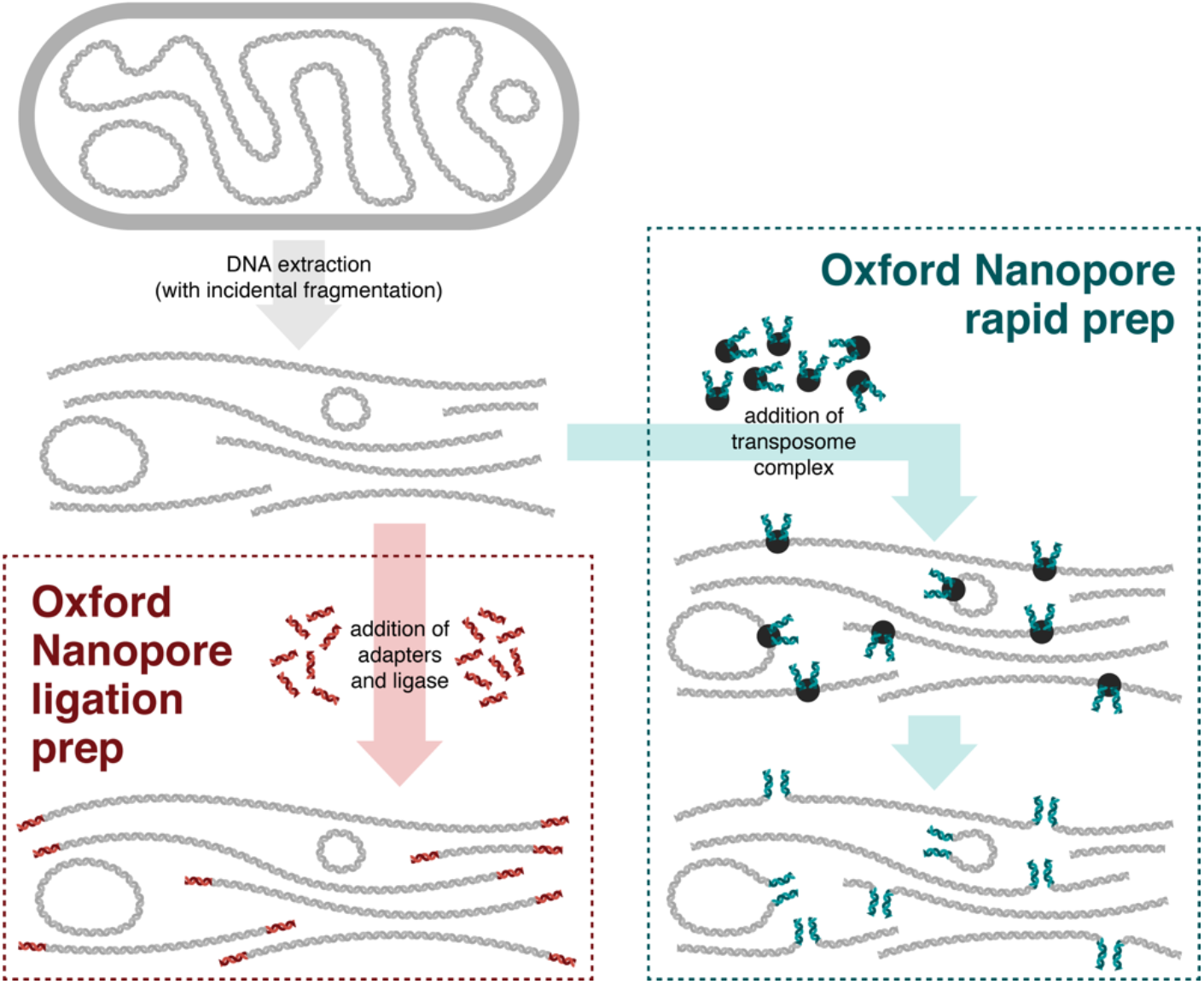
conceptual illustration of Oxford Nanopore ligation and rapid sample preparation methods. When circular DNA is extracted from a bacterial cell (top-left), incidental fragmentation of the DNA occurs. The ligation preparation (bottom-left) comprises blunt-end ligation of barcodes/adapters onto DNA molecules, so circular pieces of DNA will not receive adapters and thus remain unavailable for sequencing. The rapid preparation (right) uses a transposome enzyme to add barcodes/adapters into the middle of DNA molecules, making both linear and circular DNA available for sequencing.

The ONT rapid preparation kits (e.g. the SQK-RBK004 rapid barcoding kit) offer a potential solution. Unlike the ligation preparation approach, the rapid library approach uses a transposase enzyme to simultaneously cleave DNA and attach barcode/adapter sequences (**Figure 1**). Since rapid preparations do not rely on blunt-end ligation, they should be active on sequences that are circular such as unfragmented small plasmids. Hence, in this study we compared the performance of ONT ligation and rapid library preparations for whole-genome sequencing of bacterial genomes containing small plasmids. Specifically, we aimed to quantify plasmid read depths and determine whether ONT rapid preparations give a more accurate representation of small plasmid abundance than ONT ligation preparations. To assess this we use short-read Illumina sequencing, whose libraries are unbiased with respect to the starting length of the molecules, as the gold standard for quantifying plasmid abundance.

## Methods

### Bacterial isolates, DNA extraction and sequencing

We included seven bacterial isolates in this study (**Table 1, Figure S1A**), each containing small plasmids (identified from previous analyses) and belonging to different bacterial species: *Acinetobacter baumannii, Citrobacter koseri, Enterobacter kobei*, an unnamed *Haemophilus* species (given the placeholder name *Haemophilus sp002998595* in GTDB r95^34,35^), *Klebsiella oxytoca, Klebsiella variicola* and *Serratia marcescens*. These isolates were cultured overnight at 37°C in Luria-Bertani broth and DNA was extracted using GenFind v3 according to the manufacturer’s instructions (Beckman Coulter) (**Figure S1B**). The same DNA extract was used to sequence each isolate using three different approaches: ONT ligation, ONT rapid and Illumina (**Figure S1C**). For ONT ligation, we followed the protocol for the SQK-LSK109 ligation sequencing kit and EXP-NBD104 native barcoding expansion (Oxford Nanopore Technologies). For ONT rapid, we followed the protocol for the SQK-RBK004 rapid barcoding kit (Oxford Nanopore Technologies). All ONT libraries were sequenced on MinION R9.4.1 flow cells. For Illumina, we followed a modified Illumina DNA Prep protocol (catalogue number 20018705), whereby the reaction volumes were quartered to conserve reagents. Illumina libraries were sequenced on the NovaSeq 6000 using SP reagent kits v1.0 (300 cycles, Illumina Inc.), producing 150 bp paired-end reads with a mean insert size of 331 bp. We repeated this process (from culture to sequencing) to generate a set of technical replicates. For the first technical replicate, a refuel (with the EXP-FLP002 flow cell priming kit) was performed at the 18-hour point of the ONT runs to boost yield. However, no refuelling step was required for the second replicate. All ONT read sets were basecalled and demultiplexed using Guppy v3.6.1 (**Figure S1D–E**). Basecalled reads (FASTQ format) are available in the supplementary data repository.

**Table 1:** bacterial isolates used in this study. Each genome contained at least one large (≥20 kbp) and one small (<20 kbp) plasmid. An asterisk (*) indicates that the plasmid contains one or more antimicrobial resistance determinants.

| Isolate species and name | Large plasmids (bp) | Small plasmids (bp) |
| --- | --- | --- |
| *Acinetobacter baumannii* J9 | 145059* | 6078* |
| *Citrobacter koseri* MINF_9D | 64962* | 9294 |
| *Enterobacter kobei* MSB1_1B | 136482, 108411 | 4665, 3715, 2369 |
| *Haemophilus* M1C132_1 | 39398 | 10719, 9975*, 7392, 5675 |
| *Klebsiella oxytoca* MSB1_2C | 118161, 58472 | 4574 |
| *Klebsiella variicola* INF345 | 250980, 243620*, 31780 | 5783, 3514 |
| *Serratia marcescens* 17-147-1671 | 184477*, 161385* | 17406, 1934 |

### Reference genome assembly

In order to assign reads to their replicon of origin, we required a reference assembly for each of the seven genomes in the study. We produced these assemblies using pooled reads from all available sequencing runs (**Figure S1F**), with read QC performed by fastp v0.20.1^36^ (using default parameters) and Filtlong v0.2.0 (using a minimum read length of 1 kbp, a minimum mean quality of 80 and a minimum window quality of 60). We used Trycycler^37^ to produce consensus long-read assemblies of each isolate (using 50× subsampled read sets assembled by Flye v2.8^23^, miniasm/Minipolish v0.3/v0.1.3^38^ and Raven v1.1.10^39^), followed by polishing with Medaka v1.0.3^40^ and Pilon v1.23^41^. Technical replicates were assembled independently and the resultant assemblies compared using edlib^42^. Wherever differences were found, we examined the Illumina and Nanopore read alignments for the region in question to assess whether the difference indicated an assembly error, manually repairing such errors as appropriate. Only one genuine sequence difference was found between the two technical replicates: a 1 bp indel in the smallest plasmid of *Enterobacter kobei* MSB1_1B. Additionally, two plasmids in *Haemophilus* M1C132_1 only appeared in assemblies for the second technical replicate, as they were missing from all ONT and Illumina read sets in the first technical replicate. This suggests that they were lost during culturing in the first technical replicate. After curation, we merged the assemblies of the two technical replicates to make a single reference assembly for each of the seven isolates (**Figure S1G**), including the two plasmids that only appeared in the second technical replicate. The resultant genomes contained a total of seven chromosomes and 26 plasmid sequences (**Table 1, Figure S2**). Using a size threshold of 20 kbp, 14 of the plasmids were classified as ‘small’ and 12 as ‘large’. Seven of the plasmids (5 large and 2 small) contain one or more antimicrobial resistance genes (**Table S1**). The reference assembly sequences (FASTA format) are available in the supplementary data repository.

### Comparison of ONT library preparation methods

To aid our read-based analyses, we developed custom Python scripts (available in the supplementary data repository). We aligned each of the full ONT read sets to each sequence in the reference assemblies using minimap2 v2.17 (using the map-ont preset) and the `align_reads.py` script, which enabled alignment over the start-end junction of circular replicons. These alignments were processed with the `assign_reads.py` script, which assigned each read to a reference sequence by labelling each position of the read with the reference to which it best aligned (using the following filters: alignment identity ≥75%, alignment length ≥100 bp, mean read identity ≥80%, mean read coverage ≥50%). This script also gave a demultiplexing status to each read: correct (Guppy demultiplexing agreed with reference alignment), incorrect (Guppy demultiplexing disagreed with reference alignment), unclassified (the read was not demultiplexed by Guppy), unaligned (the read did not align to the reference sequences) or chimera (the read aligned to multiple different reference sequences). The `get_depths.py` script was used to calculate per-replicon ONT read depths by taking the mean depth across each replicon, excluding repetitive regions common to multiple replicons (identified by cross-replicon minimap2 alignments) as such repeats could make for unreliable alignments. Finally, we aligned each Illumina read set to its respective reference genome using Bowtie2 v2.3.4.1 and used the `get_depths.py` script to calculate per-replicon Illumina read depths.

Plasmid read depths were normalised to their corresponding chromosomal read depth (e.g. a plasmid with a depth twice that of the chromosomal depth had a normalised read depth of 2), providing a quantification of plasmid abundance that could be compared between read sets. Read depths were calculated separately for the two technical replicates, as plasmid copy number could differ between DNA extractions. This resulted in three normalised read depths for each plasmid in each technical replicate: ONT ligation, ONT rapid and Illumina (**Table S1**).

The on-bead tagmentation process in the Illumina DNA Prep produces libraries with a fragment size of ∼300–400 bp, which is considerably smaller than the smallest replicon in our genomes. Therefore, we assumed that unlike long-read ONT sequencing, Illumina sequencing is not significantly biased by replicon size. Since Illumina sequencing is known to have biases regarding GC content^43^, we examined this effect in our data by calculating read depth and GC content for each 1 kbp sequence window in the chromosomes of the seven study genomes (using the `depth_and_gc.py` script). By plotting the read depth (normalised to the mean depth for 50% GC windows) vs GC content, we estimated that fluctuations in GC content resulted in <10% variation of Illumina read depth (**Figure S3**). We therefore assume that our normalised Illumina read depth values are a good approximation of the true copy number of each replicon, without adjusting for GC content.

## Results and discussion

### Sequencing data characteristics

The four ONT sequencing runs (ligation run 1, rapid run 1, ligation run 2 and rapid run 2) yielded a total of 15.2 Gbp, 4.3 Gbp, 8.0 Gbp and 9.6 Gbp data, respectively. Per-barcode ONT yields ranged from 64 Mbp to 2.8 Gbp, equating to mean read depths of 10× to 1214× (see **Table S1**). Per-barcode ONT N50 read lengths ranged from 1.9 kbp to 25.8 kbp. Per-barcode Illumina yields ranged from 313 Mbp to 1.06 Gbp (40× to 232× depth, see **Table S1**). Since we used pooled ONT read sets (both ligation and rapid) to perform assemblies, all isolates had sufficient data to produce reliable reference genomes.

While previous studies have shown that ligation preparations favour read count and rapid preparations favour read length^44^, we observed no clear trends (**Table S1, Figure S4**). Many factors influence these metrics, including the quality and quantity of extracted DNA, the number of available pores on the flow cell, and the incubation times during library preparation steps. Our results show that for multiplexed bacterial whole-genome sequencing, good yields and read lengths (>100× depth and >15 kbp N50) are possible from either type of library preparation.

Library preparation type also did not seem to affect the sequence accuracy of reads. All runs had a maximum read identity of ∼98%, but two of the runs (ligation run 1 and rapid run 2) had a larger proportion of low-identity reads (22% and 23% of reads at <90% identity vs 15% and 16% for ligation run 2 and rapid run 1, respectively; see **Figure S5**). These runs suffered from degradation in translocation speed (the rate at which DNA moves through the nanopore) over the course of the run (**Figure S6**), which may partly explain their lower accuracy^45^. When this problem occurs, it can be mitigated by refuelling a run in progress using ONT’s EXP-FLP002 kit, as we did for both ligation run 1 and rapid run 1. The beneficial effect of refuelling on read accuracy was particularly notable for ligation run 1 but was negligible for rapid run 1 (**Figure S7**).

Demultiplexing accuracy was also inconsistent between preparation types, with both the best (0.39%) and worst (3.87%) demultiplexing error rates occurring in rapid runs, compared to error rates of 2.22% and 2.92% for the ligation runs (**Table S1**). During ligation preparations, barcode sequences are attached on both ends of the reads, while rapid preparations result in barcode attachment to the start of the reads only. This gives users of ligation preparations the option of running Guppy with the `--require_barcodes_both_ends` option (not used in this study) to increase demultiplexing accuracy when needed but at the expense of a much larger proportion of unclassified reads^46^.

### Chimeric read rates

The rate of chimeras (reads originating from two or more discontiguous pieces of DNA) was notably different between the two preparations: 1.41% and 0.88% chimeric reads for the two ligation runs vs 0.03% and 0.14% for the two rapid runs (**Table S1**). There are two broad categories of chimeric reads: *in silico* chimeras and ligated chimeras^47^. *In silico* chimeras occur when two separate pieces of DNA pass through a pore in quick succession such that the sequencing software mistakes them as a single read. Thus *in silico* chimeras can potentially happen for either type of preparation. Ligated chimeras occur when two separate pieces of DNA are physically joined before sequencing. Since only the ligation preparation involves ligase, ligated chimeras should be comparatively rare in rapid preparations, which may explain our observation that rapid preparations have fewer chimeric reads overall. Our results suggest that rapid preparations are preferable when chimeric reads need to be minimised. However, we note that this conclusion is derived from small sample sizes (n=2 for each preparation). Additionally, in the context of bacterial whole-genome assembly, we have previously shown that chimeric read rates of up to ∼5% (i.e. exceeding those of our sequencing runs) do not impact assembly quality^24^.

### Small plasmid abundance

For each assembled plasmid, we calculated a normalised depth ratio: its normalised ONT read depth (i.e. ONT plasmid depth relative to the chromosome) divided by its normalised Illumina read depth (i.e. Illumina plasmid depth relative to the chromosome). Ratios greater than one indicate the plasmid was overrepresented in ONT reads relative to Illumina reads, and ratios less than one indicate the plasmid was underrepresented. **Figure 2** shows the relationship between normalised depth ratios and plasmid size for ligation and rapid ONT preparations.

**Figure 2:**
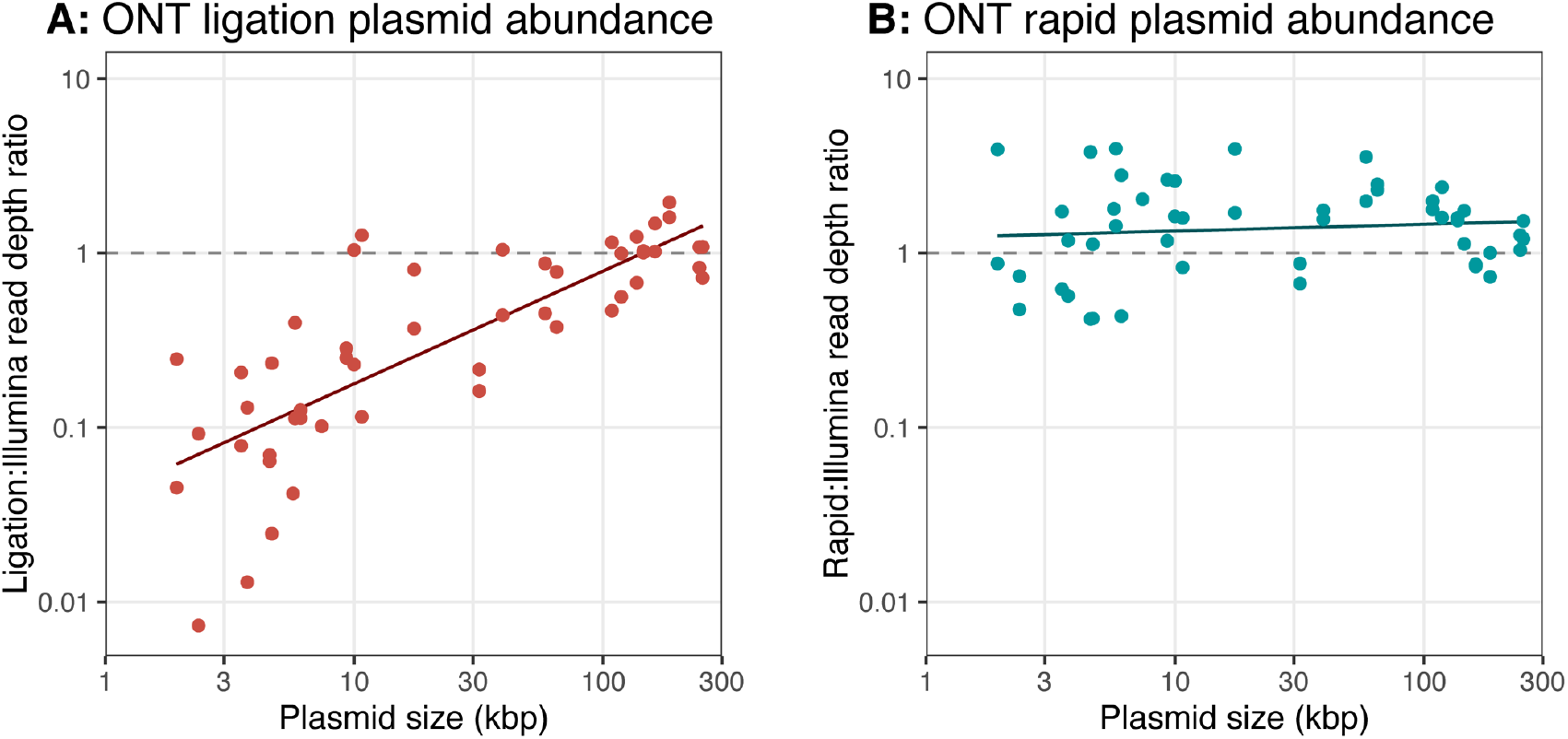
plasmid abundance resulting from A) ligation and B) rapid ONT library preparation methods. Each point in the plots represents one plasmid. The read depth ratio is the normalised ONT read depth divided by the normalised Illumina read depth. The dashed lines at ratio=1 indicate perfect agreement of plasmid depths between ONT and Illumina data. Points above the dashed lines indicate plasmids that are overrepresented in ONT reads, while points below the dashed lines indicate plasmids that are underrepresented in ONT reads. The solid lines indicate least-squares linear regressions of the log-transformed data. For ONT ligation reads (A), small plasmids are systematically underrepresented relative to Illumina reads. For ONT rapid reads (B), plasmid size has no clear effect, and depths for both small (<20 kbp) and large plasmids (≥20 kbp) are in good agreement with Illumina reads.

For ONT ligation reads, there was a clear relationship between the normalised depth ratio and plasmid size (*p*=5.8×10^−12^, *R*^2^=0.63, 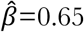, from a linear regression performed on log-transformed data, *H*_0_ of β=0) (**Figure 2A**). Specifically, ONT ligation reads tended to underrepresent small plasmids (<20 kbp), for which the mean normalised depth ratio was 25% (i.e. small plasmids produced ∼4× fewer ONT ligation reads than one would expect based on Illumina read depths). The most extreme case was for the 2.4 kbp plasmid in the second technical replicate of *E. kobei* MSB1_1B, which had a normalised depth ratio of <1%, i.e. it was underrepresented by a factor of more than 100. ONT rapid read depths, however, showed no relationship with plasmid size (*p*=0.47, *R*^2^=0.011, 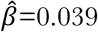, using the same statistical test, **Figure 2B**).

These results suggest that both ONT ligation and ONT rapid preparations provide an accurate representation of abundance for very large plasmids (>100 kbp), but only the ONT rapid preparation accurately represents abundance across the full spectrum of plasmid sizes. While these conclusions are based on the assumption that Illumina reads accurately represent plasmid abundance, this is supported by the fact that Illumina and ONT rapid plasmid read depths are in good agreement.

The inherent bias against small plasmids during ligation-based library preparations could result in the exclusion of small plasmids from genome assemblies, particularly when performing long-read-only assembly (as opposed to hybrid assembly where Illumina reads are also available) and assembling read sets of modest depth. For example, if a genome containing a four-copies-per-cell small plasmid was sequenced with a chromosomal read depth of 25×, the plasmid sequence should be present at ∼100× depth. However, if that small plasmid was underrepresented by a factor of 30 (a realistic possibility based on our data, see **Figure 2A**), it might only be sequenced at ∼3× depth, making it unlikely to appear in an assembly for that genome.

### Conclusions

When faced with the choice of ligation or rapid ONT preparations, researchers must weigh their respective advantages. Ligation kits are versatile, can give greater yields and allow for multiplexing of more samples. Rapid kits are faster, require fewer additional resources and equipment, and can provide longer read lengths (if optimised for in DNA preparation^44^). Our study reveals two more advantages of the rapid kits: lower chimeric read rates and better recovery of small plasmids. Rapid preparations are therefore more likely to produce long-read-only assemblies which include all of a genome’s replicons, small plasmids included.

## Author statements

### Authors and contributors

Conceptualization: RRW, KLW, KEH. Methodology: RRW. Software: RRW. Formal analysis: RRW, KLW. Investigation: RRW, LMJ. Resources: RRW, LMJ. Data curation: RRW, LMJ. Writing–original draft: RRW. Writing–review and editing: RRW, LMJ, KLW, KEH. Visualization: RRW. Supervision: KEH, KLW. Project administration: KEH, KLW. Funding acquisition: KEH.

### Conflicts of interest

The authors declare that there are no conflicts of interest.

### Funding information

This work was supported by the Bill & Melinda Gates Foundation, Seattle (grant number OPP1175797) and an Australian Government Research Training Program Scholarship. KEH is supported by a Senior Medical Research Fellowship from the Viertel Foundation of Victoria. The funders had no role in study design, data collection and analysis, decision to publish, or preparation of the manuscript.

## Notes

https://github.com/rrwick/Small-plasmid-Nanopore

### Competing Interest Statement

The authors have declared no competing interest.

## References

1. Land M, Hauser L, Jun S-R, Nookaew I, Leuze MR, Ahn T-H, et al. Insights from 20 years of bacterial genome sequencing. Functional & Integrative Genomics. 2015;15(2):141–61. doi:10.1007/s10142-015-0433-4.

2. Bobay LM, Ochman H. The evolution of bacterial genome architecture. Frontiers in Genetics. 2017;8(MAY):1–6. doi:10.3389/fgene.2017.00072.

3. Baker S, Hardy J, Sanderson KE, Quail M, Goodhead I, Kingsley RA, et al. A novel linear plasmid mediates flagellar variation in *Salmonella* Typhi. PLoS Pathogens. 2007;3(5):0605–10. doi:10.1371/journal.ppat.0030059.

4. Arredondo-Alonso S, Top J, McNally A, Puranen S, Pesonen M, Pensar J, et al. Plasmids shaped the recent emergence of the major nosocomial pathogen *Enterococcus faecium*. mBio. 2020;11(1):1– 17. doi:10.1128/mBio.03284-19.

5. Ciok A, Dziewit L, Grzesiak J, Budzik K, Gorniak D, Zdanowski MK, et al. Identification of miniature plasmids in psychrophilic Arctic bacteria of the genus *Variovorax*. FEMS Microbiology Ecology. 2016;92(4):1–9. doi:10.1093/femsec/fiw043.

6. Rozwandowicz M, Brouwer MSM, Fischer J, Wagenaar JA, Gonzalez-Zorn B, Guerra B, et al. Plasmids carrying antimicrobial resistance genes in *Enterobacteriaceae*. Journal of Antimicrobial Chemotherapy. 2018;73(5):1121–37. doi:10.1093/jac/dkx488.

7. Shintani M, Sanchez ZK, Kimbara K. Genomics of microbial plasmids: Classification and identification based on replication and transfer systems and host taxonomy. Frontiers in Microbiology. 2015;6(MAR):1–16. doi:10.3389/fmicb.2015.00242.

8. Smalla K, Jechalke S, Top EM. Plasmid detection, characterization and ecology. Plasmids. 2015;3(1):445–58. doi:10.1128/9781555818982.ch23.

9. San Millan A, Escudero JA, Catalan A, Nieto S, Farelo F, Gibert M, et al. β-Lactam resistance in *Haemophilus parasuis* is mediated by plasmid pB1000 bearing blaROB-1. Antimicrobial Agents and Chemotherapy. 2007;51(6):2260–4. doi:10.1128/AAC.00242-07.

10. Anantham S, Hall RM. pCERC1, a small, globally disseminated plasmid carrying the *dfrA14* cassette in the *strA* Gene of the *sul2-strA-strB* Gene Cluster. Microbial Drug Resistance. 2012;18(4):364–71. doi:10.1089/mdr.2012.0008.

11. Lanza VF, de Toro M, Garcillán-Barcia MP, Mora A, Blanco J, Coque TM, et al. Plasmid flux in *Escherichia coli* ST131 sublineages, analyzed by plasmid constellation network (PLACNET), a new method for plasmid reconstruction from whole genome sequences. PLOS Genetics. 2014;10(12). doi:10.1371/journal.pgen.1004766.

12. Wick RR, Judd LM, Gorrie CL, Holt KE. Unicycler: Resolving bacterial genome assemblies from short and long sequencing reads. PLOS Computational Biology. 2017;13(6):e1005595. doi:10.1371/journal.pcbi.1005595.

13. Arredondo-Alonso S, Willems RJ, van Schaik W, Schürch AC. On the (im)possibility of reconstructing plasmids from whole-genome short-read sequencing data. Microbial Genomics. 2017;3(10). doi:10.1099/mgen.0.000128.

14. Conlan S, Thomas PJ, Deming C, Park M, Lau AF, Dekker JP, et al. Single-molecule sequencing to track plasmid diversity of hospital-associated carbapenemase-producing *Enterobacteriaceae*. Science Translational Medicine. 2014;6(254):254ra126.

15. Weingarten RA, Johnson RC, Conlan S, Ramsburg AM, Dekker JP, Lau AF, et al. Genomic analysis of hospital plumbing reveals diverse reservoir of bacterial plasmids conferring carbapenem resistance. mBio. 2018;9(1):1–16. doi:10.1128/mBio.02011-17.

16. Loman NJ, Quick J, Simpson JT. A complete bacterial genome assembled *de novo* using only nanopore sequencing data. Nature Methods. 2015;12(8):733–5. doi:10.1038/nmeth.3444.

17. Koren S, Phillippy AM. One chromosome, one contig: Complete microbial genomes from long-read sequencing and assembly. Current Opinion in Microbiology. 2015;23:110–20. doi:10.1016/j.mib.2014.11.014.

18. Wick RR, Judd LM, Gorrie CL, Holt KE. Completing bacterial genome assemblies with multiplex MinION sequencing. Microbial Genomics. 2017;3(10):1–7. doi:10.1099/mgen.0.000132.

19. Taylor TL, Volkening JD, DeJesus E, Simmons M, Dimitrov KM, Tillman GE, et al. Rapid, multiplexed, whole genome and plasmid sequencing of foodborne pathogens using long-read nanopore technology. Scientific Reports. 2019;9(1):1–11. doi:10.1038/s41598-019-52424-x.

20. Elliott I, Batty EM, Ming D, Robinson MT, Nawtaisong P, De Cesare M, et al. Oxford nanopore MinION sequencing enables rapid whole genome assembly of rickettsia typhi in a resource-limited setting. American Journal of Tropical Medicine and Hygiene. 2020;102(2):408–14. doi:10.4269/ajtmh.19-0383.

21. Antipov D, Korobeynikov A, McLean JS, Pevzner PA. HybridSPAdes: An algorithm for hybrid assembly of short and long reads. Bioinformatics. 2016;32(7):1009–15. doi:10.1093/bioinformatics/btv688.

22. Koren S, Walenz BP, Berlin K, Miller JR, Phillippy AM. Canu: Scalable and accurate long-read assembly via adaptive *k*-mer weighting and repeat separation. Genome Research. 2017;27:722– 36. doi:10.1101/gr.215087.116.

23. Kolmogorov M, Yuan J, Lin Y, Pevzner PA. Assembly of long, error-prone reads using repeat graphs. Nature Biotechnology. 2019;37(5):540–6. doi:10.1038/s41587-019-0072-8.

24. Wick RR, Holt KE. Benchmarking of long-read assemblers for prokaryote whole genome sequencing. F1000Research. 2019;8(2138). doi:10.12688/f1000research.21782.1.

25. Lam MMC, Wick RR, Wyres KL, Gorrie CL, Judd LM, Jenney AWJ, et al. Genetic diversity, mobilisation and spread of the yersiniabactin-encoding mobile element ICE*Kp* in *Klebsiella pneumoniae* populations. Microbial Genomics. 2018;4(9). doi:10.1099/mgen.0.000196.

26. Lam MMC, Wyres KL, Judd LM, Wick RR, Jenney A, Brisse S, et al. Tracking key virulence loci encoding aerobactin and salmochelin siderophore synthesis in *Klebsiella pneumoniae*. Genome Medicine. 2018;10(1):1–15. doi:10.1186/s13073-018-0587-5.

27. Wyres KL, Hawkey J, Hetland MAK, Fostervold A, Wick RR, Judd LM, et al. Emergence and rapid global dissemination of CTX-M-15-associated *Klebsiella pneumoniae* strain ST307. Journal of Antimicrobial Chemotherapy. 2019;74(3):577–81. doi:10.1093/jac/dky492.

28. Lam MMC, Wyres KL, Wick RR, Judd LM, Fostervold A, Holt KE, et al. Convergence of virulence and MDR in a single plasmid vector in MDR *Klebsiella pneumoniae* ST15. Journal of Antimicrobial Chemotherapy. 2019;74(5):1218–22. doi:10.1093/jac/dkz028.

29. Wyres KL, Wick RR, Judd LM, Froumine R, Tokolyi A, Gorrie CL, et al. Distinct evolutionary dynamics of horizontal gene transfer in drug resistant and virulent clones of *Klebsiella pneumoniae*. PLOS Genetics. 2019;15(4):1–25. doi:10.1371/journal.pgen.1008114.

30. Wyres KL, Nguyen TNT, Lam MMC, Judd LM, van Vinh Chau N, Dance DAB, et al. Genomic surveillance for hypervirulence and multi-drug resistance in invasive *Klebsiella pneumoniae* from South and Southeast Asia. Genome Medicine. 2020;12(1):11. doi:10.1186/s13073-019-0706-y.

31. Jorgensen JH, Ferraro MJ. Antimicrobial susceptibility testing: A review of general principles and contemporary practices. Clinical Infectious Diseases. 2009;49(11):1749–55. doi:10.1086/647952.

32. Klingström T, Bongcam-Rudloff E, Pettersson OV. A comprehensive model of DNA fragmentation for the preservation of High Molecular Weight DNA. bioRxiv. 2018. doi:10.1101/254276.

33. Ou S, Liu J, Chougule KM, Fungtammasan A, Seetharam AS, Stein JC, et al. Effect of sequence depth and length in long-read assembly of the maize inbred NC358. Nature Communications. 2020;11(1):1–10. doi:10.1038/s41467-020-16037-7.

34. Chaumeil PA, Mussig AJ, Hugenholtz P, Parks DH. GTDB-Tk: A toolkit to classify genomes with the genome taxonomy database. Bioinformatics. 2020;36(6):1925–7. doi:10.1093/bioinformatics/btz848.

35. Parks DH, Chuvochina M, Chaumeil PA, Rinke C, Mussig AJ, Hugenholtz P. A complete domain-to-species taxonomy for Bacteria and Archaea. Nature Biotechnology. 2020. doi:10.1038/s41587-020-0501-8.

36. Chen S, Zhou Y, Chen Y, Gu J. Fastp: An ultra-fast all-in-one FASTQ preprocessor. Bioinformatics. 2018;34(17):i884–90. doi:10.1093/bioinformatics/bty560.

37. Wick RR, Holt KE. Trycycler. GitHub. 2020. url:github.com/rrwick/Trycycler.

38. Li H. Minimap and miniasm: Fast mapping and de novo assembly for noisy long sequences. Bioinformatics. 2016;32(14):2103–10. doi:10.1093/bioinformatics/btw152.

39. Vaser R, Mile Š, Šikić M. Raven: A *de novo* genome assembler for long reads. bioRxiv. 2020. doi:10.1101/2020.08.07.242461.

40. Wright C, Wykes M. Medaka. GitHub. 2020. url:github.com/nanoporetech/medaka.

41. Walker BJ, Abeel T, Shea T, Priest M, Abouelliel A, Sakthikumar S, et al. Pilon: An integrated tool for comprehensive microbial variant detection and genome assembly improvement. PLOS ONE. 2014;9(11). doi:10.1371/journal.pone.0112963.

42. Šošić M, Šikić M. Edlib: A C/C ++ library for fast, exact sequence alignment using edit distance. Bioinformatics. 2017;33(9):1394–5. doi:10.1093/bioinformatics/btw753.

43. Aird D, Ross MG, Chen W-S, Danielsson M, Fennell T, Russ C, et al. Characterizing and measuring bias in sequence data. Genome biology. 2012;02(5):1. doi:10.1186/gb-2011-12-2-r18.

44. Jain M, Koren S, Quick J, Rand AC, Sasani TA, Tyson JR, et al. Nanopore sequencing and assembly of a human genome with ultra-long reads. Nature Biotechnology. 2018;36:338–345. doi:10.1038/nbt.4060.

45. Amarasinghe SL, Su S, Dong X, Zappia L, Ritchie ME, Gouil Q. Opportunities and challenges in long-read sequencing data analysis. Genome Biology. 2020;21(1):1–16. doi:10.1186/s13059-020-1935-5.

46. Wick RR, Judd LM, Holt KE. Deepbinner: Demultiplexing barcoded Oxford Nanopore reads with deep convolutional neural networks. PLOS Computational Biology. 2018;14(11):e1006583. doi:10.1371/journal.pcbi.1006583.

47. White R, Pellefigues C, Ronchese F, Lamiable O, Eccles D. Investigation of chimeric reads using the MinION. F1000Research. 2017;6(May):631. doi:10.12688/f1000research.11547.1.

